# Developing mammary terminal duct lobular units have a dynamic mucosal and stromal immune microenvironment

**DOI:** 10.1101/2020.11.05.369843

**Authors:** Dorottya Nagy, Clare M. C. Gillis, Katie Davies, Abigail L. Fowden, Paul Rees, John W. Wills, Katherine Hughes

## Abstract

The human breast and ovine mammary gland undergo a striking degree of postnatal development, leading to formation of terminal duct lobular units (TDLUs). In this study we interrogated aspects of sheep TDLU growth to increase understanding of ovine mammogenesis and as a model for the study of breast development. Mammary epithelial proliferation is significantly higher in lambs less than two months old than in peri-pubertal animals. Ki67 expression is polarized to the leading edge of the developing TDLUs. Intraepithelial ductal macrophages exhibit striking periodicity and significantly increased density in lambs approaching puberty. Stromal macrophages are more abundant centrally than peripherally. The developing ovine mammary gland is infiltrated by intraepithelial and stromal T lymphocytes that are significantly more numerous in older lambs. In the stroma, hotspots of Ki67 expression colocalize with large aggregates of lymphocytes and macrophages. Multifocally these aggregates exhibit distinct organization consistent with tertiary lymphoid structures. The lamb mammary gland thus exhibits a dynamic mucosal and stromal immune microenvironment and, as such, constitutes a valuable model system that provides new insights into postnatal breast development.

**Summary statement:** Development of terminal duct lobular units in the sheep mammary gland involves distinct growth phases and macrophage and lymphocyte fluxes. Tertiary lymphoid structures are present subjacent to the mucosal epithelium.

## Introduction

The mammary gland undergoes a dramatic degree of postnatal growth, developing from a rudimentary branched structure at birth to an arborizing bilayered ductal network in the nulliparous adult.

Macrophages are key players in the direction of murine mammary ductal growth (Brady et al., 2016) and there is increasing recognition of a spectrum of mammary macrophage subsets (Wilson et al., 2020b). Mammary macrophages may be derived from the foetal liver and yolk sac and additionally infiltrate postnatally (Jäppinen et al., 2019). Depletion experiments have demonstrated the dependence of mammary postnatal development on macrophages (Gouon-Evans et al., 2000) and that alveolar bud formation and ductal epithelial proliferation are reduced in their absence (Chua et al., 2010). Stat5 is activated in mammary macrophages during development, and mice with macrophages that have conditional deletion of Stat5 exhibit perturbed development (Brady et al., 2017). Cells expressing MHCII are closely associated with murine mammary ducts (Hitchcock et al., 2020, Dawson et al., 2020), and macrophages envelop the pubertal terminal end buds (Stewart et al., 2019). The atypical chemokine receptor ACKR2, which scavenges CC-chemokines, has been implicated in macrophage recruitment during mammary development (Wilson et al., 2017, Wilson et al., 2020a). Intriguingly, macrophage depletion of virgin mice also influences the mammary stromal extracellular matrix composition, highlighting the importance of macrophages in both the epithelial and stromal compartments (Wang et al., 2020).

CD4+ and CD8+ lymphocytes have also been identified in the murine mammary gland (Plaks et al., 2015, Betts et al., 2018). As T-cell receptor alpha deficient mice exhibit enhanced ductal outgrowths, it is postulated that T-lymphocytes may act in a negative regulatory manner (Plaks et al., 2015). Similarly, lymphocytes are present in the human breast (Howard and Gusterson, 2000, Degnim et al., 2014) although little is known about their developmental role.

An understanding of postnatal pre-pregnancy breast development in humans is critical to interrogation of the pathogenesis of breast diseases (Osin et al., 1998). Whilst mouse models of mammary development are highly tractable and extremely valuable, they have inherent limitations and caution has been recommended in the extrapolation of results of murine developmental studies directly to humans (Gusterson and Stein, 2012). Potentially pertinent given the complex interactions between cellular compartments, mammary epithelial cells in the breast are surrounded by fibrous connective tissue whereas the murine mammary stroma is adipose-rich (Hovey et al., 1999). By contrast, the ruminant mammary gland exhibits a strikingly similar micro-anatomical arrangement of terminal duct lobular units (TDLUs) and fibrous stroma to the human breast (Hovey et al., 1999, Hughes and Watson, 2018a). We and others have therefore suggested that it represents a valuable adjunctive model of the breast TDLU (Rowson et al., 2012, Hughes, 2020) although further interrogation of the utility of this model is required.

Sheep are frequently used as a model species in foetal development studies (Morrison et al., 2018) and also constitute a globally valuable production animal species. However, there are currently a number of knowledge gaps concerning the biology of ruminant mammogenesis (Davis, 2017) and a better understanding of ovine-specific mammary development is required to underpin attempts to breed animals for improved milk production efficiency and reduced susceptibility to mastitis.

Given that studying ovine mammary development will offer new insights relevant to breast development, and that there is a pressing need for species-specific data regarding udder development in the pre-pregnancy ewe, we sought to capitalise on the availability of new technologies to study postnatal mammary development in this species. We utilised deep learning image analysis to define phases of growth in ovine mammary TDLU development and employed 2-dimensional and deep 3-dimensional (3D) imaging approaches to interrogate and quantify the presence of macrophages, lymphocytes and tertiary lymphoid structures within the gland during development.

## Results and Discussion

### Mammary epithelial proliferation is significantly higher in younger lambs than in those approaching puberty, with proliferation focused at the leading edge of the advancing TDLUs

Preclinical models of tumourigenesis do not always portray the heterogeneity of human disease (Cassidy et al., 2015), and this limitation may also apply to developmental studies where a relatively homogeneous population of rodents, maintained in controlled conditions, may not recapitulate the diversity of the progression of breast development noted in humans (Howard and Gusterson, 2000). For this study we therefore selected a heterogeneous population of pre- and peri-pubertal lambs of differing breeds, maintained in different husbandry systems. This population of lambs exhibit developing TDLUs supported by intra- and interlobular stroma (Figs. S1, S2), very similar to the breast, and in contrast to the murine mammary gland (Hovey et al., 1999).

To assess nulliparous ovine mammary growth dynamics, we performed immunohistochemical staining (IHC) for Ki67 to delineate actively cycling cells. There is significant up-regulation of Ki67 expression in lambs less than 2 months old compared to peri-pubertal lambs aged 5-9.5 months old (Fig. 1A-D). This finding is similar to that recorded in a small study of infant breasts where epithelial Ki67 positivity was not detected after 25 days of age (Osin et al., 1998). It also builds upon a historic study using dried fat-free tissue weights to assess mammary growth that suggested that ovine allometric mammary growth occurred at 3-4 months old, prior to puberty. Notably, that analysis was somewhat limited in scope, with only Romney and Romney-cross animals examined and no animals older than 5 months old included in the pre-pregnancy group (Anderson, 1975).

**Figure 1.**
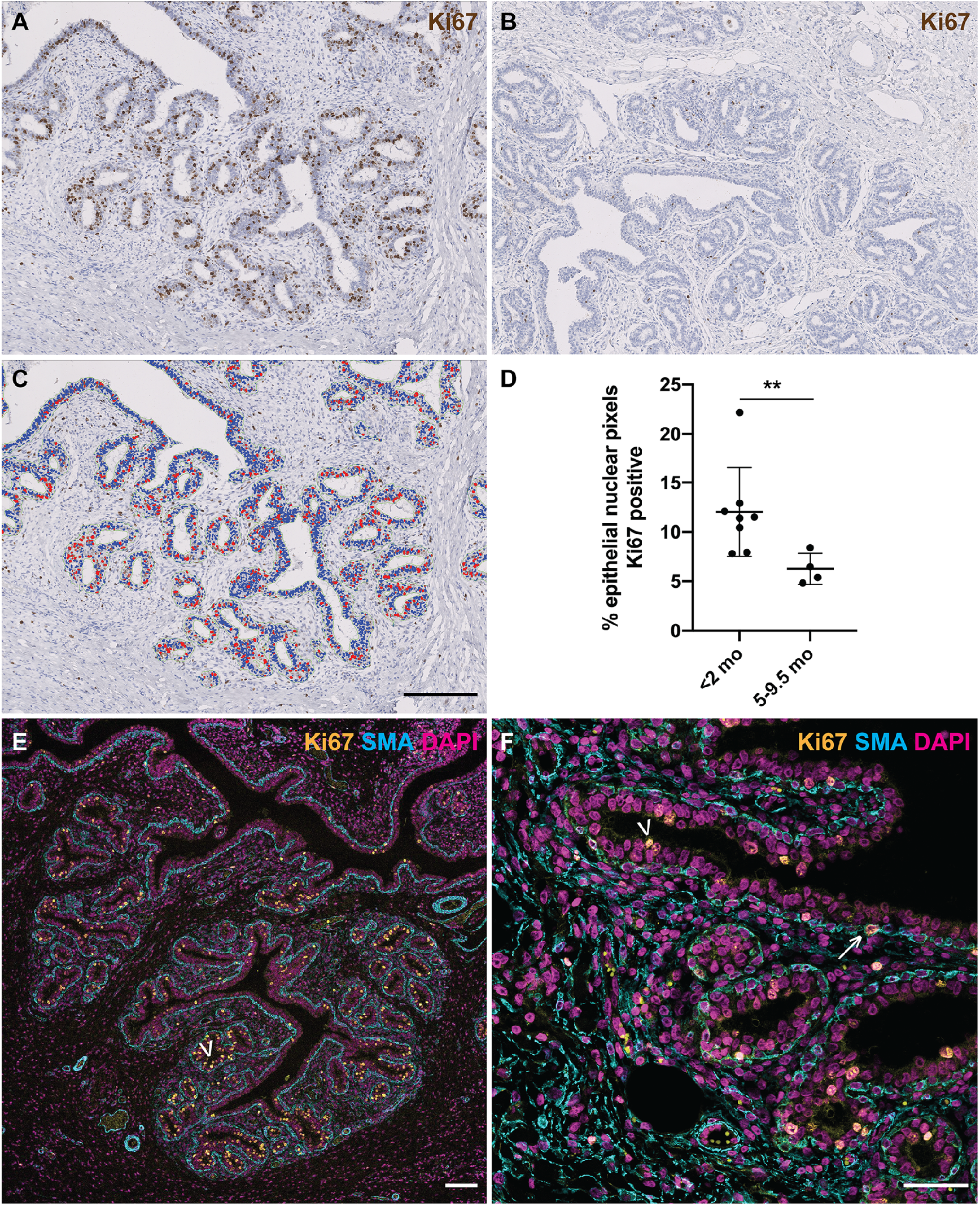
Mammary epithelial proliferation is significantly higher in younger lambs than in those approaching puberty. (A-C) IHC for Ki67 in mammary gland from lambs < 2 mo (A) and 5-9.5 mo (B) and accompanying mask derived using an algorithm detecting intra-epithelial Ki67 positive events (C). (D) Scatter plot demonstrating significantly higher levels of epithelial nuclear Ki67 positivity in younger lambs. Dots represent individual lambs. Bars represent mean +/- standard deviation. ** p < 0.01. (E-F) IF for Ki67 (gold), α-SMA (cyan) and DNA (DAPI; magenta) demonstrating that the majority of Ki67 positive nuclei are in the luminal epithelial layer (arrowheads), with rare Ki67 positive nuclei in myoepithelial cells (arrow). (E) 1 do lamb. (F) 9.5 mo lamb. do, days old; mo, months old. Images are representative of a minimum of three biological repeats. All IHC images have haematoxylin counterstain. Scale bar = 200 μm (A-C); 100 μm (E); 50 μm (F).

In the present study, immunofluorescence staining (IF) demonstrates that although the majority of epithelial proliferation is luminal, myoepithelial (basal) cells occasionally express Ki67 (Fig. 1E,F). This highlights similarities with the breast, where sporadic proliferating myoepithelial cells have been noted in normal breast parenchyma of women aged 30 to 68 years, using samples where biopsies or mass removal has included normal tissue (Bankfalvi et al., 2004). There has been a relative paucity of focus on myoepithelial proliferation within the developing breast or mammary gland prior to pregnancy. During lactation, myoepithelial cells contract to deform alveoli, facilitating milk release in response to oxytocin stimulation (Stevenson et al., 2020). Our identification of proliferation within the myoepithelial compartment pre-pregnancy suggests that studying basal epithelial replication during this period may provide new insights into udder development relevant to lactation efficiency.

Having observed that pre-pregnancy ovine mammary epithelial proliferation is not temporally uniform, we wished to interrogate the spatial distribution of Ki67-positive epithelial events. Spatial statistical analyses (Getis-Ord GI*) reveal distinct polarization of epithelial proliferation towards the advancing tips of the developing TDLUs (Fig. 2A-D), echoing non-quantified description of non-random localization of Ki67 expression in the infant breast (Osin et al., 1998). This finding further underlines the utility of the lamb mammary gland as a model of breast development. Interestingly, qualitative descriptions of a similar phenomenon of Ki67 polarization have also been made in rats, where Ki67 positivity is focused in the terminal end buds (Hvid et al., 2012).

**Figure 2.**
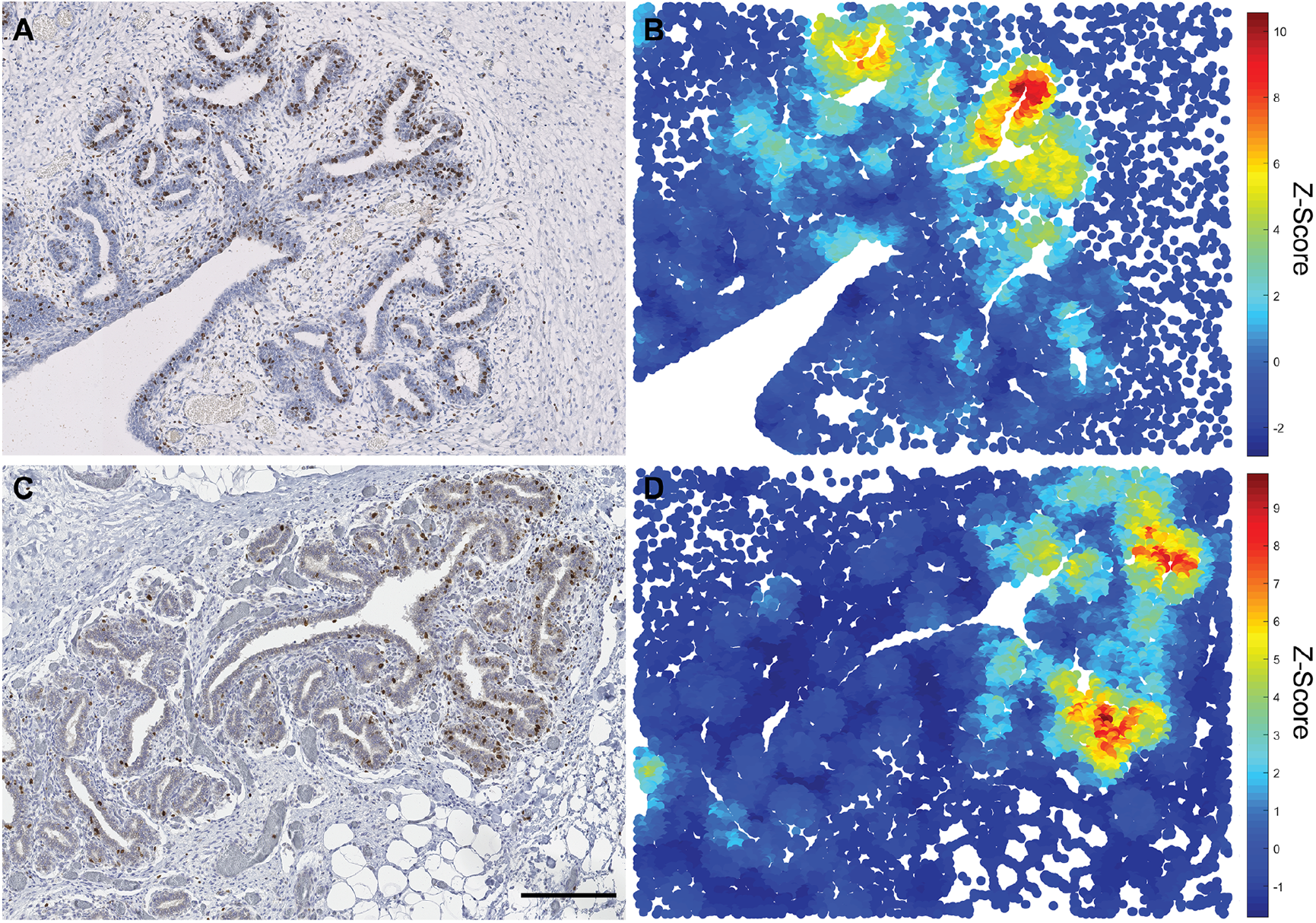
The developing lamb mammary gland exhibits polarity of Ki67 epithelial expression with Ki67 expression focused at the leading edge of the advancing TDLUs. IHC for Ki67 (A, C) and accompanying Getis-Ord (G-O) statistical analyses (B, D) demonstrating regions with significant spatial congregation of intraepithelial Ki67+ cells (scale *(d)* parameter = 250 px). Mammary gland from lambs < 2 mo (A, B) and 5-9.5 mo (C, D). (A, C) Haematoxylin counterstain. Scale bar = 200 μm. (B, D). Results are representative of four biological repeats (two lambs in each age group).

### Macrophages exhibit spatial and temporal dynamics within the pre-pregnancy TDLU

Having established that the ovine mammary gland exhibits a distinct growth phase during pre-pubertal mammary development, we wished to compare the spatial and temporal distribution of macrophages during development pre-pregnancy. The macrophage marker ionized calcium binding adaptor molecule 1 (IBA1) is expressed by macrophages and microglia and is involved in macrophage membrane ruffling (Ohsawa et al., 2000). We have previously utilized this marker to detect ovine mammary macrophages (Hardwick et al., 2020). In the present study, using IBA1 IHC to identify macrophages, we noted distinct periodicity of intraepithelial macrophages both with ducts and ductules (Fig. 3A,B) similar to that reported in mice (Stewart et al., 2019, Dawson et al., 2020). Importantly, we identified a previously unrecognized variation in ductular macrophage density, with a significantly reduced inter-macrophage distance in ducts examined from peri-pubertal animals (Fig. 3C). This increased ductular macrophage density may suggest enhanced immune surveillance in animals approaching puberty, or a reorganization of macrophage distribution following the pulse of growth associated with pre-pubertal development.

**Figure 3.**
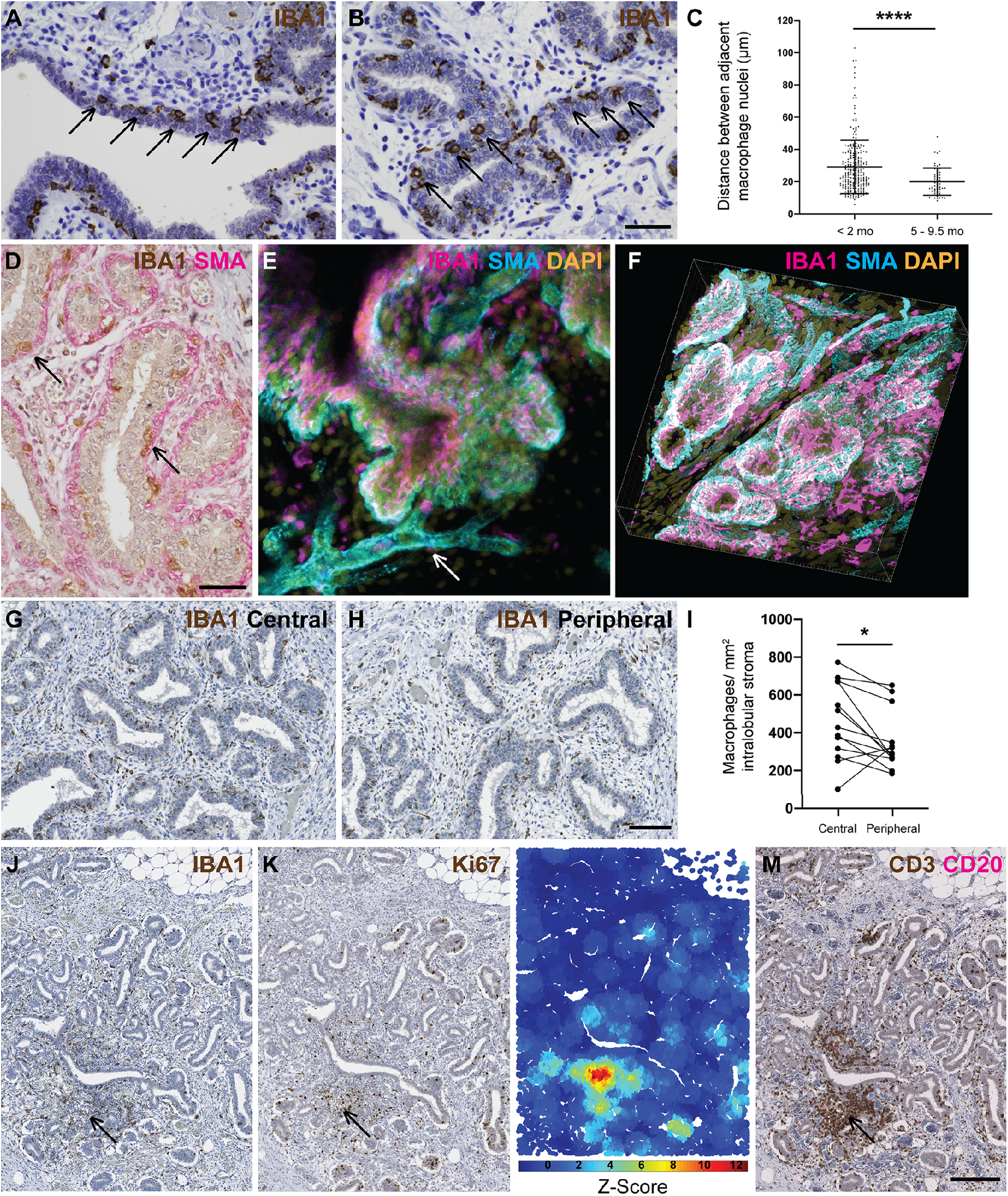
Mammary macrophages exhibit spatial and temporal dynamics. (A-B) IHC for IBA1 reveals macrophage periodicity (arrows) in ducts (A) and ductules (B). (C) Scatter plot demonstrating significantly reduced inter-macrophage distance in lambs aged 5-9.5 months. Dots represent inter-macrophage distances from 13 individual lambs. Bars represent mean +/- standard deviation. **** p < 0.0001. (D) IHC for IBA1 (brown) and alpha smooth muscle actin (SMA; pink). Arrows indicate macrophages. (E-F) 3D confocal microscopy of optically cleared ovine mammary tissue with IF for IBA1 (magenta) SMA (cyan) and DNA (Hoechst; gold). Images represent 3D maximum intensity projections. Arrow indicates blood vessel. (G-H) IHC for IBA1 in central (G) and peripheral (H) foci. (I) Scatter plot demonstrating significantly reduced macrophage abundance in peripheral compared to central foci. Dots represent average macrophage density for individual lambs. * p < 0.05. (J-M) Serial sections demonstrating IHC for IBA1 (J) Ki67 (K) and CD3 (brown) and CD20 (pink) (M) with accompanying G-O plot for Ki67 (L) (scale *(d)* parameter = 250 px). Arrow indicates co-localization of stromal macrophages, a Ki67 hotspot, and a CD3+ lymphocyte aggregate. Images are representative of a minimum of three biological repeats. All IHC images have haematoxylin counterstain. Scale bar = 40 μm (A,B,D); 100 μm (G-H); 200 μm (J,K,M).

Within the developing TDLU, macrophages are intercalated within the ductal epithelial bilayer similar to the arrangement reported in the mouse (Fig. 3D-F; Movie 1) (Dawson et al., 2020). The TDLU-associated ductal macrophages form a largely contiguous layer sandwiched between the luminal and basal epithelial cells. We hypothesise that during development pre-pregnancy this complex of macrophages is likely to fulfil an immune surveillance function, commensurate with a proposed ability to sample the epithelium through movement of cellular processes (Dawson et al., 2020) and underlining the concept of the mammary ductular microenvironment as a mucosal immune system (Betts et al., 2018).

In addition to an abundant intraepithelial macrophage population, frequent macrophages are present in the ovine intralobular stroma encasing the developing TDLUs. Interestingly, these stromal macrophages are more numerous in central foci than in peripheral locations (Fig. 3G-I). This may point to stromal macrophage abundance surrounding the developing ruminant gland cistern (Fig. S2), likely reflecting an important role in immune regulation of the mammary microenvironment. However, murine stromal macrophages derived from adult mice have differing gene expression profiles compared to ductal macrophages (Dawson et al., 2020). It is thus probable that stromal macrophages also have other functions. A recent study focusing on mammary stromal macrophages has delineated a homeostatic role for this population, with Lyve-1 expressing stromal macrophages associated with areas of hyaluronan enrichment in both mice and humans. Mice in which macrophages were depleted exhibited increased levels of hyaluronan within the stromal adipose (Wang et al., 2020). It is therefore possible that the abundance of stromal macrophages that we have noted in the central portion of the developing ruminant TDLU may reflect mesenchymal remodelling as the gland cistern develops.

Although stromal macrophages usually exhibit a relatively regular distribution (Fig. 3G,H), we noted multifocal stromal foci in which there are more dense aggregates of IBA1 positive macrophages admixed with lymphocytes (Fig. 3J). Intriguingly, these correspond to hotspots of Ki67 expression (Fig. 3K,L). The aggregates are predominantly composed of CD3-expressing T lymphocytes, with variable numbers of CD20-expressing B lymphocytes (Fig. 3M). This prompted us to further investigate lymphocyte distribution within the developing ovine TDLUs.

### Epithelial and stromal T lymphocytes are more abundant in older lambs than in neonates, and stromal lymphocytes multifocally form tertiary lymphoid structures

Both intraepithelial and stromal CD3+ T lymphocytes are significantly more abundant in older lambs than in neonates (Fig. 4A-D). CD4+ T helper 1 lymphocytes have previously been identified as negative regulators of mammary development and so it is tempting to speculatively associate the abundance of intraepithelial T lymphocytes in older lambs with the observed decrease in epithelial proliferation within the TDLU in this age group. The mammary immune system has been likened to a classical mucosal immune system (Betts et al., 2018) and the presence of mammary intraepithelial lymphocytes is reminiscent of other mucosal surfaces such as the intestinal epithelium, where intraepithelial lymphocytes are common (Cheroutre et al., 2011). Notably, the CD3+ T lymphocytes in mammary intraepithelial foci frequently exhibit a similar spatial niche to intraepithelial macrophages, intercalated between the luminal and basal epithelial layers (Fig. 3D-F; Fig. 4E). We and others have previously described mammary intraepithelial lymphocytes in rabbits, mice and humans respectively (Hughes and Watson, 2018b, Plaks et al., 2015, Degnim et al., 2014) and so it seems likely that this distribution is common to many species.

**Figure 4.**
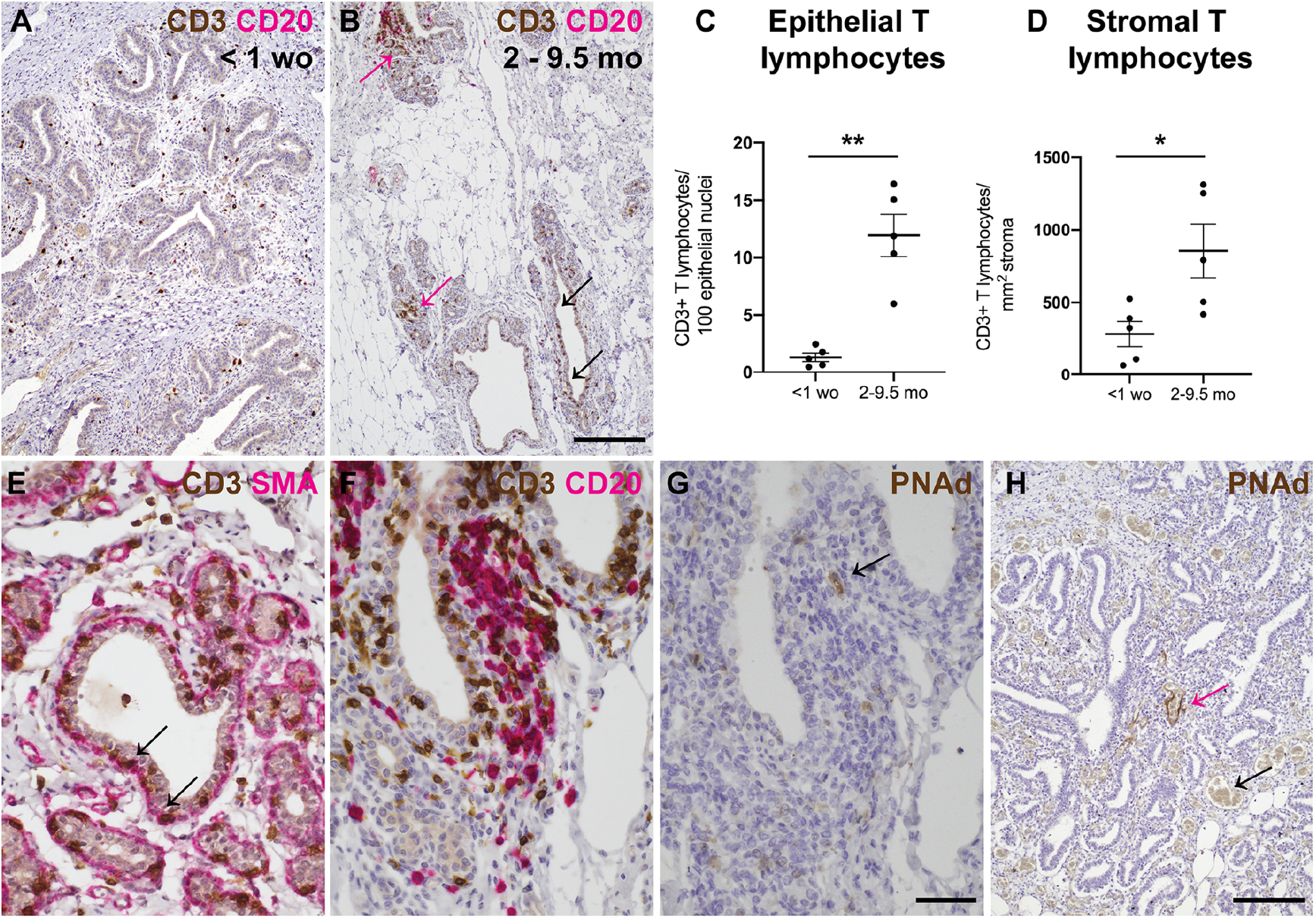
T lymphocytes are more abundant in older lambs than in neonates, and tertiary lymphoid structures are multifocally present. (A-B) IHC for CD3 (brown) and CD20 (pink) reveals more abundant intraepithelial (black arrows) and stromal (pink arrows) T lymphocytes in older lambs. (C-D) Scatter plots demonstrating significantly increased numbers of epithelial (C) and stromal (D) T lymphocytes in older lambs. Dots represent CD3+ lymphocyte densities from individual lambs. Bars represent mean +/- standard deviation. * p < 0.05; ** p < 0.01. (E) IHC for CD3 (brown) and SMA (pink). Arrows indicate intraepithelial lymphocytes. (F) IHC for CD3 (brown) and CD20 (pink). An aggregate of lymphocytes in a subepithelial focus exhibits a central zone of B lymphocytes surrounded by T lymphocytes. (G-H) IHC for PNAd. (G) Serial section of (F). Arrow indicates high endothelial venule within the aggregate of lymphocytes depicted in (F). (H) Pink arrow indicates PNAd-positive blood vessel amidst lymphocytic infiltrate. Black arrow indicates adjacent negative internal control blood vessel, demonstrating specificity of staining. Images are representative of a minimum of three biological repeats. All IHC images have haematoxylin counterstain. Scale bar = 200 μm (A,B); 40 μm (E-G); 200 μm (H).

Finally, we noted that some stromal aggregates of T and B lymphocytes exhibit distinct arrangement with central foci of B lymphocytes surrounded by a more peripheral of T lymphocytes. High endothelial venules, denoted by expression of peripheral node addressin (PNAd), are detectable within these aggregates (Fig. 4F,G) and the groupings exhibit characteristics of tertiary lymphoid structures (TLS). TLS are aggregates of lymphocytes possessing distinct architectural arrangement, similar to secondary lymphoid organs, which may arise in foci of chronic inflammation, or secondary to autoimmune processes or neoplasia (Pipi et al., 2018). In our study, the density of TLS did not differ significantly between neonatal and older lambs, although less tissue area per lamb was available for examination from the neonatal lambs and this may have reduced the likelihood of detecting a TLS (Fig. S3).

The finding that TLS are present subjacent to the mammary mucosal epithelium is particularly important given that the pre- and peri-pubertal animals studied had never lactated and never exhibited evidence of clinical or subclinical mastitis. Indeed, mastitis would be extremely rare in this age group. Therefore the occurrence of TLS may indicate that mammary pre-lactational subclinical pathogen challenge is common in lambs.

These observations suggest that the formation of TLS immediately subjacent to mammary ducts in pre-pregnancy animals may constitute a hitherto unrecognised component of the mammary gland’s mucosal immune system. It seems likely that these structures may form in response to antigenic stimulation reflecting the contiguity between the mammary epithelium and the epidermis (Betts et al., 2018). Corroborating this finding, we noted that small calibre blood vessels located in foci of mixed T and B lymphocyte aggregates, lacking the zonal organization of TLS, also mutifocally and selectively express endothelial PNAd (Fig. 4H). Such vascular expression of PNAd has been suggested to be associated with ‘immature’ foci in which less organised lymphocyte groupings are in the process of forming TLS (Ager, 2017). Thus the formation of TLS is likely an active ongoing process in nulliparous lambs.

One benefit of the present study is that much larger tissue areas are typically available for analysis from ovine subjects compared to those likely available from infant breast tissue, or from normal tissue present adjacent to surgically removed breast lesions. Therefore it is possible that TLS are a feature of the mammary mucosal immune system of other species but these structures may be rarely detectable in the samples available to researchers.

Our work demonstrates that ovine developing mammary TDLUs have a dynamic mucosal and stromal immune microenvironment. We provide valuable new data on the growth phases and macrophage and lymphocyte fluxes occurring prior to gestation and document that TLS do not solely arise as a result of mastitis (Restucci et al., 2019) but rather are a naturally occurring component of the lamb mammary immune microenvironment. We also demonstrate a number of similarities between the ovine mammary gland and human breast. The lamb mammary gland thus constitutes a valuable model system that provides new insights into postnatal breast development.

## Materials and Methods

### Animals

Mammary tissue was collected for this study from two separate sources. Mammary tissue was collected from female sheep aged less than one year that were submitted to the diagnostic veterinary anatomic pathology service of the Department of Veterinary Medicine, University of Cambridge. Additionally, mammary tissue was obtained post mortem from 2 day old – 9.5 months old Welsh mountain ewes studied for other research purposes (Davies et al., 2020) and euthanased under the Animals (Scientific Procedures) Act 1986. The Ethics and Welfare Committee of the Department of Veterinary Medicine, University of Cambridge, approved the study plan relating to the use of ovine post mortem material for the study of mammary gland biology (reference: CR223). The non-regulated scientific use of post mortem mammary tissue collected from research animals was approved by the Named Veterinary Surgeon of the University of Cambridge. Together, sheep from these two sources comprised a range of hill breeds and crosses, aged between 0 days and 9.5 months.

In all cases, macro- and microscopic post mortem examination of mammary tissue was conducted by a single American board-certified veterinary pathologist and no tissues with macro-or microscopic mammary pathology were included in the study.

### Histology

Mammary tissue was fixed in 10% neutral-buffered formalin for approximately 72 hours. Tissues were processed and tissue sections were cut at five microns. These were stained with haematoxylin and eosin.

### Immunohistochemistry and immunofluorescence

Antibodies utilised for immunohistochemical (IHC) and immunofluorescence (IF) staining are detailed in Supplementary Table 1. IHC followed a routine protocol using a PT link antigen retrieval module and high pH antigen retrieval solution (both Dako Pathology/Agilent Technologies, Stockport, UK). Primary and secondary antibodies were incubated for 1 hour at room temperature. For dual IHC staining, an ImmPRESS™ Duet Double Staining Polymer Kit (Vector laboratories, Peterborough, UK) was utilised. Negative control slides were prepared using isotype- and species-matched immunoglobulins or secondary antibody only.

IF also followed a routine protocol. Antigen retrieval was carried out using a PT link antigen retrieval module and high pH antigen retrieval solution as detailed above. Primary antibodies were incubated overnight at 4°C and secondary antibodies were incubated for 1 hour at room temperature. Nuclei were stained with DAPI (10.9 μM) (Sigma-Aldrich/Merck Life Science UK Limited, Gillingham, UK). Slides were mounted using Vectashield® Vibrance™ Antifade mounting medium (catalogue H-1700; Vector laboratories, Peterborough, UK). Imaging was performed using either a Leica TCS SP8 or a Zeiss LSM780 confocal microscope.

### Tissue clearing and deep 3D imaging

Tissues were optically cleared using the CUBIC protocol as previously described (Susaki et al., 2014, Lloyd-Lewis et al., 2016) with minor modifications as detailed below. Ovine mammary tissue was cut into slices approximately 10 mm thick and was fixed for 6-8 hours in 10% neutral-buffered formalin. Tissue was then sufficiently firm to be cut into smaller pieces, on average 5×8×2 mm. Tissue pieces were subsequently immersed in CUBIC reagent 1A for 4 days at 37 °C with gentle rocking. The CUBIC reagent 1A solution was replaced daily. Samples were blocked in blocking buffer comprising normal goat serum [10% (volume per volume)] and Triton X-100 [0.5% (weight per volume)] in PBS. Samples were blocked overnight at 4 °C with gentle agitation. Tissue samples were incubated with primary antibodies diluted in blocking buffer for 4 days at 4 °C with gentle agitation. The samples were then washed at room temperature with gentle rocking in PBS containing Triton X-100 (0.1% (weight per weight)). Secondary antibodies were also prepared in blocking buffer and tissue samples were incubated in these for 2 days at 4 °C, with gentle rocking. Following thorough washing as described above, samples were incubated with DAPI (10.9 μM) (Sigma-Aldrich/Merck Life Science UK Limited, Gillingham, UK) for a minimum of 1 hour at room temperature prior to further washing and immersion in CUBIC reagent 2 for at least 2 days at 37 °C with gentle rocking. Negative control tissue was prepared by omitting the primary antibody and using the secondary antibody only. Cleared and stained tissue fragments were imaged in Ibidi 35 mm glass bottom dishes (catalogue 81218-200; ibidi GmbH, Gräfelfing, Germany) using a Leica TCS SP8 confocal microscope. 3D data were visualised using ImarisViewer (Oxford Instruments, UK. Imaris Viewer: a free 3-D/4-D microscopy image viewer. https://imaris.oxinst.com/imaris-viewer Accessed 03/11/2020) and Vaa3D (Peng et al., 2014) software.

### Slide scanning

Slides IHC stained for Ki67, IBA1, and CD3/CD20 were scanned at 40× using a NanoZoomer 2.0RS, C10730, (Hamamatsu Photonics, Hamamatsu City, Japan). Scanned sections were analysed with viewing software (NDP.view2, Hamamatsu Photonics).

### Computational analyses

#### Ki67: Deep Learning Image Analysis

130 image-fields (DAB Ki67^+^ detection/haematoxylin counterstain) each covering 1.5 mm^2^ (6322 x 4581 pixels) were collected from slide scans across animals in RGB tiff format. Images were normalised across the haematoxylin / DAB colour-components using the Macenko approach (Macenko et al., 2009). Fourteen image fields were used to train the deep learning models. Firstly, a two class, semantic pixel classification network (DeepLabV3+ on a pre-trained ResNet18 backbone with output stride eight (He et al., 2016, Chen et al., 2018) was trained to provide a binary mask of ‘epithelium’ or ‘background/other’ classes. Input images were passed to the network as patches (2000/image) with dimensions 256, 256, 3 (x, y, channels) and augmented by random x/y reflection and rotation. The network was trained for 150 epochs using a batch size of eight with zero-centre normalisation under stochastic gradient descent using class-weighted cross-entropy loss. The initial learn rate was 0.001 with a drop factor every ten epochs of 0.3, a momentum of 0.9 and L2 regularisation 0.05. Patches were shuffled every epoch.

To segment Ki67^+^ and Ki67^-^ nuclei, a three-class (‘Ki67^+^ nuclei’, ‘Ki67^-^ nuclei’, ‘background/other’) Unet model (Ronneberger et al., 2015) was trained - again using data from fourteen, Macenko-normalised image fields. Patches (2000/image) were passed to the network with dimensions 256, 256, 3 (x, y, channels) and simple augmentation by random x/y reflection and rotation. The Unet model utilised an encoder depth of four layers with 64 filters in the first layer. The network used complete, up-convolutional expansion to yield images identically sized to the input layer. Training lasted for fifty epochs, using batch size of eight with zero-centre normalisation under stochastic gradient descent utilising cross-entropy loss. The initial learn rate was 0.05, dropping every ten epochs by 0.1 under momentum 0.9 and L2 regularisation 0.0001.

Models were trained using MATLAB R2020 and the Deep Learning Toolbox. The trained models, test data alongside all training hyper-parameters and final layer-weightings are available for download at BioStudies database (http://www.ebi.ac.uk/biostudies) under accession number S-BSST528. Both models were tested against entirely unseen data (the other 116 fields) and the results validated using boundary overlays and manual image-review by an American board-certified veterinary pathologist. The ratio (pixel area) of Ki67^+^ to Ki67^-^ nuclei in the epithelium of each image-field was calculated using the epithelial segmentation mask from the DeepLabV3+ResNet18 model to mask the Unet segmentations for each nuclear phenotype.

#### Ki67: Getis-Ord Spatial Analyses

Per-nuclei intensity and spatial location data were extracted using CellProfiler (Carpenter et al., 2006) as described in previous work (Wills et al., 2020). Statistically significant, spatial ‘congregations’ of Ki67^+^ nuclei relative to what would be expected by random chance were identified using the Getis-Ord GI* statistical approach (Ord and Getis, 1995). Ki67^+^ and Ki67^-^ nuclear objects segmented by the Unet model were used to define the centroid position for both nuclear phenotypes in an image-field. The spatial concentration of values *x_j_* for *j* values within a distance *d* of the value *x_i_* were then defined. To do this, the ratio 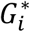 was defined as:

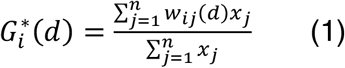

here, *w_ij_(d)* defines the numerator contribution of the ratio depending on the distance *d.* For example, using *w_ij_(d)* = 1, if *d_ij_*< *d* else; *w_ij_(d)* = 0 if *d_ij_*> *d*. From here, the Getis-Ord statistic is given by:

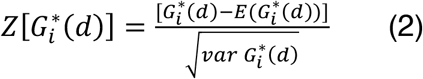

Where, 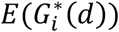 represents the expected fraction of items within *d*, assuming a completely random distribution calculated as:

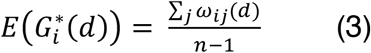

The value 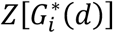 now describes the difference in the fraction of values within the distance *d* from location *i* from what would be expected by random chance relative to the standard deviation. Here, we discretise each image field into a grid and value *x_i_* is defined as the number of nuclei of a certain phenotype in the grid position *i* (Wills et al., 2020).

### Manual histopathological assessments

#### Assessment of macrophage periodicity

Macrophage periodicity was defined on IBA1 IHC stained sections as a segment of at least 4 evenly spaced intraepithelial macrophages. Spacing between macrophages was measured from the central aspect of the macrophage nucleus to the central aspect of the next macrophage nucleus using the NDP.view2 software. The centre of the cell was inferred in instances where the nucleus was not perfectly sectioned but where there was a strong impression of the nuclear position. Measurements were made parallel to the epithelium. Groups of macrophages were excluded unless they constituted a very tightly clustered small group of less than 3 macrophages in a region of clear periodicity.

#### Sampling for stromal macrophage and T lymphocyte counts

Using NDP.view2 slide viewing software, eight count boxes (400×230 μm; 4 per central or peripheral location for macrophages) were placed per slide, separately for macrophage and T lymphocyte quantification, at 1.3x magnification where only ductal structure, but not staining, was discernible, to prevent placement bias while maximising the epithelium sampled. Boxes in any fields with slide cutting artefacts or scanning focus artefacts were repositioned. For the macrophage analysis, selected fields were classified as ‘peripheral’ if sampling the edge of ductal/lobular epithelium, advancing into surrounding adipose tissue, or ‘central’ if mammary parenchyma was adjacent to the sampled area.

#### Cell quantification for stromal macrophage and T lymphocyte counts

Cells with >50% of their nucleus within the count box, or if equivocal, those along the top and right edges, were counted. A macrophage was counted as an area of IBA-1 expression that was at least 50% of the average luminal epithelial cell nucleus in that count box. ‘Stromal macrophage’ count was normalised to intralobular stromal area, determined using the NDP.view2 freehand annotation tool. ‘Epithelial T lymphocytes’ had >50% of their cytoplasmic perimeter contacting the basement membrane, with counts normalised per 100 luminal epithelial nuclei in the count box. ‘Stromal T lymphocyte’ count was normalised to total stromal area within the count box, determined using the NDP.view2 freehand annotation tool.

#### Lymphocyte aggregate qualitative description and density

TLS were defined as a discrete B lymphocyte aggregates with a distinct adjacent T lymphocyte area following previously published work (Buisseret et al., 2017). TLS were counted by two independent observers (DN and KH). Where there was a discrepancy between the counts made by the two investigators, count results from both investigators were reviewed and the final decision on count was made by the American board-certified veterinary pathologist having reviewed the identified structures. The area of mammary tissue analysed for each lamb was determined as above, using the NDP.view2 freehand annotation tool.

### Statistical Analysis

Data was recorded using Excel and analysed with GraphPad Prism 8.4.3. Immune cell counts were compared using Student’s unpaired two-tailed T-test or paired two-tailed T-test as appropriate (H0= no difference between populations).

## Acknowledgements

The authors gratefully acknowledge the excellent technical expertise of Debbie Sabin in the preparation of histology sections and unstained tissue sections. Some confocal microscopy images were acquired using equipment at the Cambridge Advanced Imaging Centre (CAIC) and the authors thank members of the CAIC for their advice and support. The Ethics and Welfare Committee of the Department of Veterinary Medicine, University of Cambridge, reviewed the study plan relating to the use of ruminant tissue collected following post mortem examination for the study of mammary gland biology (reference: CR223) and the work of this committee is gratefully recognised. The data detailed in this manuscript were presented in part at the 2020 Winter Meeting of the Pathological Society of Great Britain & Ireland (presentation: 21 January 2020) and the 2020 American College of Veterinary Pathologists Annual Meeting (presentation: 30 October 2020).

## Author contributions

Conceptualization: K.H.; Design/Methodology: P.R., J.W.W., K.H.; Validation: D.N., C.M.C.G., J.W.W., K.H. Formal analysis: J.W.W., K.H.; Investigation: D.N., C.M.C.G., P.R., J.W.W, K.H.; Resources: K. D., A.L.F., K.H.; Writing - original draft: K.H.; Writing - review & editing: D.N., A.L.F., J.W.W., K.H.; Supervision: J.W.W., K.H.; Funding acquisition: K.H.

## Data availability

Trained deep learning models, test data alongside all training hyper-parameters, and final layer-weightings are available for download at BioStudies database (http://www.ebi.ac.uk/biostudies) under accession number S-BSST528.

## Supplementary data

**Figure S1.**
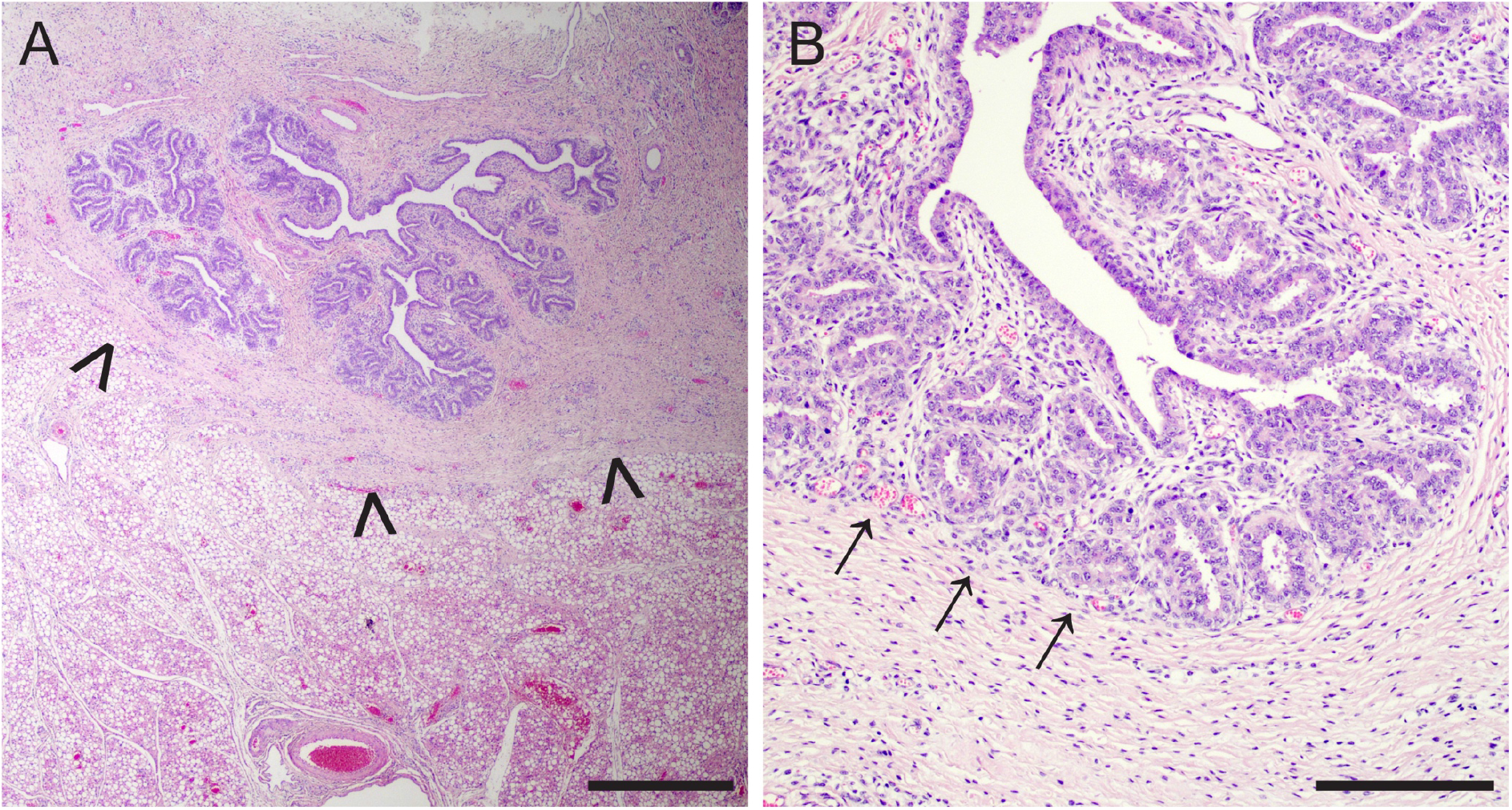
Developing ovine terminal duct lobular units (TDLUs) are supported by intra- and interlobular stroma. (A) At birth, the ovine mammary gland comprises a rudimentary structure composed of ducts and developing TDLUs. Arrowheads indicate the boundary with the deeper mammary fat pad, and correspond to the boundary indicated by arrowheads on Figure S2A. (B) The lamb mammary gland exhibits distinct intra- and interlobular stroma. Arrows indicate boundary between intra- and interlobular stroma. Haematoxylin and eosin stain. Bar: 800 microns (A); 200 microns (B). Images are representative of at least three biological repeats from lambs less than one week old.

**Figure S2.**
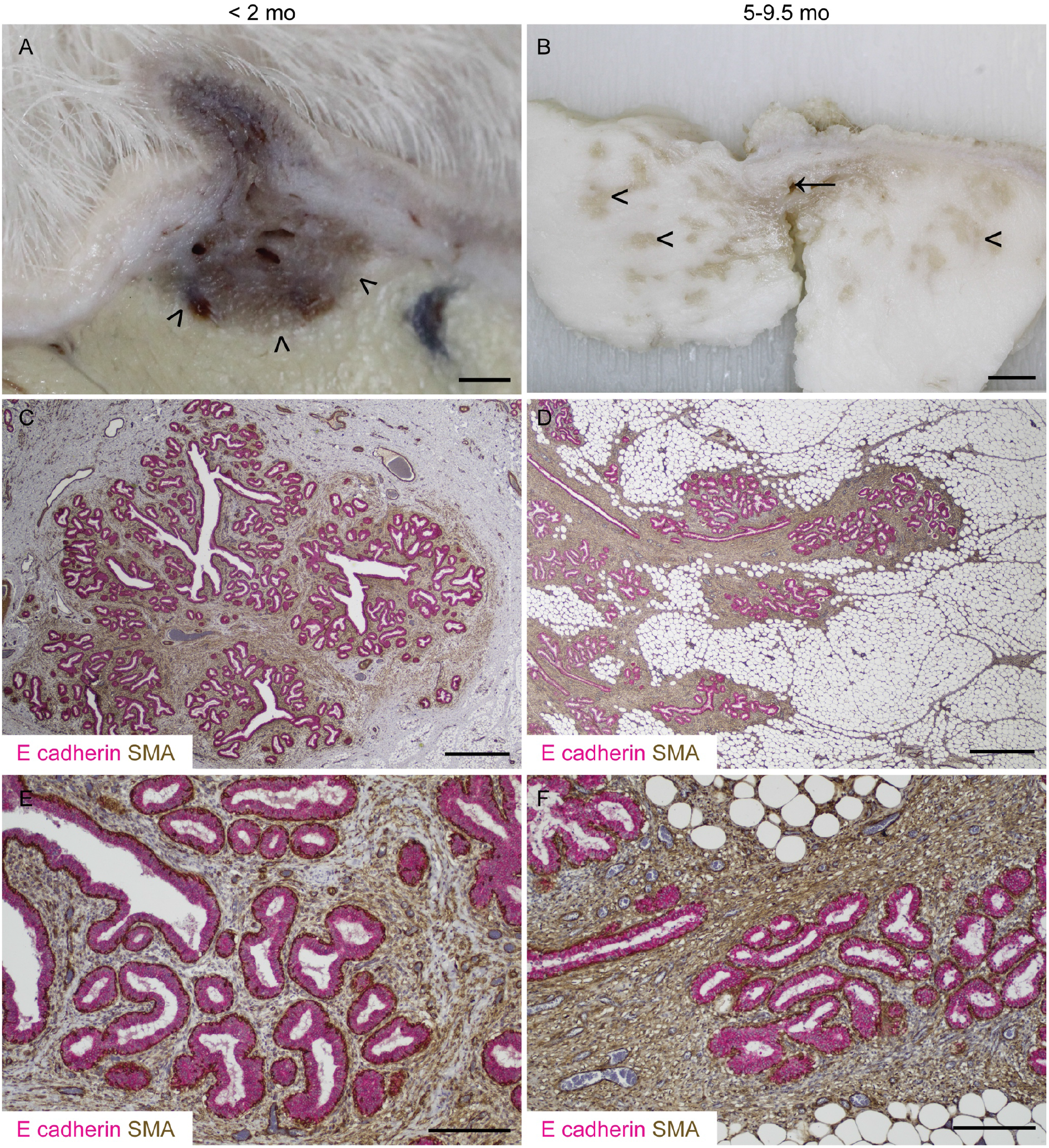
Lamb terminal duct lobular units (TDLUs) advance into the mammary fat pad during postnatal development. (A, B) Sub-gross images of fixed mammary tissue. Arrowheads indicate the developing mammary TDLUs infiltrating the mammary fat pad. Arrow indicates rudimentary gland cistern. (C-F) Immunohistochemical staining for E-cadherin (magenta) & alpha-smooth muscle actin (SMA; brown). Haematoxylin counterstain. Bar: 1.5 mm (A); 5 mm (B); 800 microns (C, D); 200 microns (E, F). Images are representative of at least three biological repeats.

**Figure S3.**
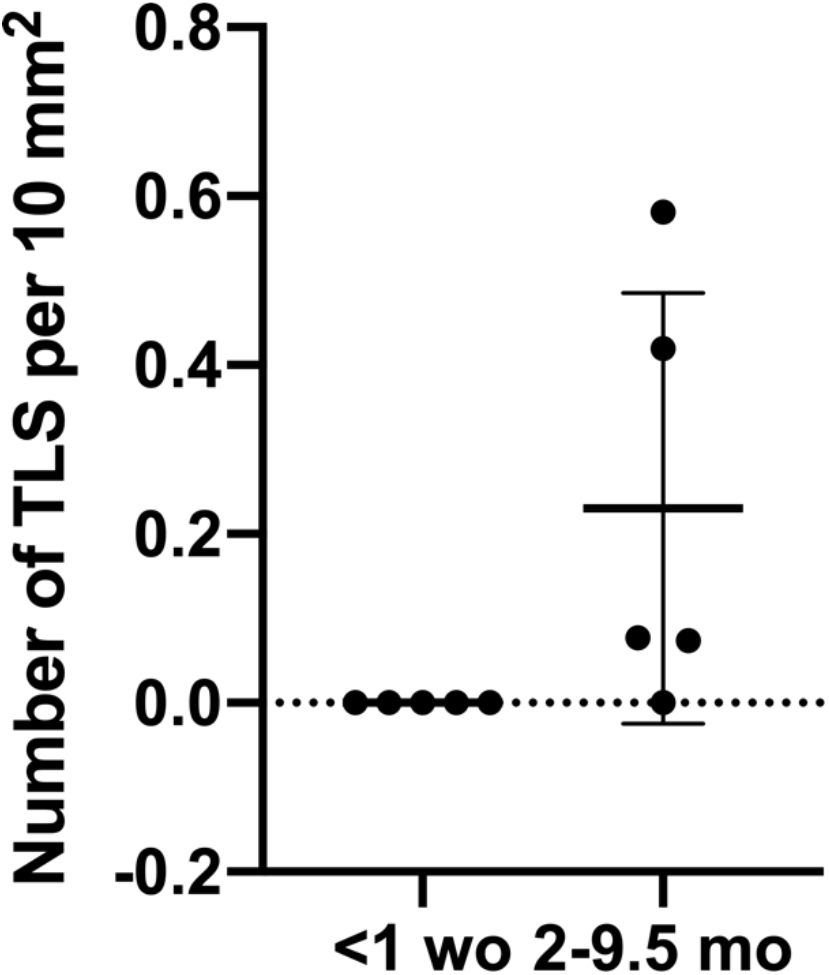
The density of tertiary lymphoid structures (TLS) does not differ significantly between neonatal and older lambs. Scatter plot demonstrating density of TLS in lambs less than one week old (< 1 wo) and aged 2-9.5 months (2-9.5 mo). Dots represent individual lambs. Bars represent mean +/- standard deviation.

**Movie 1: Three-dimensional rendering demonstrating the intimate association between myoepithelial cells (SMA; grey) and macrophages (IBA1; magenta) in CUBIC-cleared developing lamb mammary TDLUs.**

**Table S1.** Antibodies employed for immunohistochemistry, immunofluorescence, and CUBIC.

| Target | Application<br>(IHC,<br>immunohistochemistry<br>; IF,<br>immunofluorescence;<br>CUBIC, 3D tissue<br>clearing) | Species<br>and clone | Dilution | Manufacturer | Catalogue<br>number |
| --- | --- | --- | --- | --- | --- |
| **Primary antibodies** |  |  |  |  |  |
| Alpha smooth muscle actin | IF; dual colour IHC | Rabbit monoclonal [EPR5368] | 1:2000 | Abcam | Ab124964 |
| Alpha smooth muscle actin | CUBIC; dual colour IHC | Mouse monoclonal anti-human 1A4 | 1:100 (CUBIC)<br>1:400 (dual colour IHC) | Dako/Agilent | M0851 |
| CD3 | Dual colour IHC | Mouse monoclonal anti-human clone F7.2.38 | 1:250 | Dako/Agilent | M7254 |
| CD20 | Dual colour IHC | Rabbit polyclonal | 1:800 | Thermo Fisher Scientific | RB-9013-P1 |
| E-cadherin | Dual colour IHC | Rabbit monoclonal | 1:400 | Cell Signaling Technology | #3195 |
| IBA1 | IHC; dual colour IHC | Mouse monoclonal , clone 20A12.1 | 1:800 | Millipore | MABN92 |
| IBA1 | CUBIC | Rabbit monoclonal [EPR16588] | 1:400 | Abcam | Ab178846 |
| Ki67 | IHC; IF | Mouse monoclonal anti-human clone MIB-1 | 1:100 | Dako/Agilent | M7240 |
| PNA<sup>d</sup> | IHC | Rat monoclonal anti-mouse/ human clone MECA-79 | 1:100 | BioLegend | 120802 |
| **Secondary antibodies** |  |  |  |  |  |
| Mouse IgG, Alexa Fluor Plus 488 | IF | Goat | 1:500 | Thermo Fisher Scientific | A32723 |
| Mouse IgG, Alexa Fluor 568 | IF | Goat | 1:500 | Thermo Fisher Scientific | A11031 |
| Rabbit IgG, Alexa Fluor Plus 488 | CUBIC | Goat | 1:500 | Thermo Fisher Scientific | A32731 |
| Rabbit IgG, Alexa Fluor Plus 647 | CUBIC | Goat | 1:500 | Thermo Fisher Scientific | A32733 |
| Rat IgG, peroxidase labelled | IHC | Goat | 1:400 | Vector laboratories | PI-9400 |

